# Hungry catfish – effect of prey availability on movement dynamics of a top predator

**DOI:** 10.1101/2024.08.30.610378

**Authors:** Milan Říha, Rubén Rabaneda-Bueno, Lukáš Vejřík, Ivan Jarić, Marie Prchalová, Marek Šmejkal, Martin Čech, Vladislav Draštík, Petr Blabolil, Michaela Holubová, Tomáš Jůza, Karl Ø. Gjelland, Zuzana Sajdlová, Luboš Kočvara, Michal Tušer, Jiří Peterka

## Abstract

1. The availability and spatial distribution of prey determines the energetic costs of predators foraging and largely drives their space use, activity level and foraging timing. Consequently, predators in ecosystems with different prey availability and distribution should adjust their movement patterns to optimise their foraging success in order to maximise efficiency.
2. The aim of this study was to investigate the ability of a top predator, European catfish (*Silurus glanis*), to adapt its space use, temporal activity and diet patterns in environments with varying prey density and distribution, and to examine the consequences for the catfish’s body growth.
3. Catfish activity, tracked with high-resolution acoustic telemetry positioning systems deployed in two oligotrophic lakes with limited fish prey and one eutrophic reservoir with abundant fish prey, showed clear differences in their spatial and foraging behaviour. In oligotrophic, prey-poor lakes, catfish showed larger space use, altered diurnal activity patterns, reduced use of the open water, increased variability in inter- and intra-individual space use and had a more diverse diet compared to their conspecifics in the eutrophic, prey-rich reservoir. However, despite these behavioural alterations the reduced food availability in prey-poor lakes ultimately led to a reduced catfish growth, as these behavioural changes could not fully compensate for the lower prey fish abundance.
4. The study showed that top predators can combine different behavioural mechanisms and individual strategies to adapt to different ecological contexts. The observed ability of catfish to optimize activity, space use and diet highlights the evolutionary need to maximise foraging efficiency and overall fitness in different habitats, giving catfish a significant competitive advantage. Such competitive abilities are prerequisites for the actual catfish success as an invasive species within a Europe.

## Introduction

Predator movements and activities directly affect their energy budget, growth and fitness (Papastamatiou et al., 2024). Their movement patterns are dynamic and strongly influenced by environmental factors, local conditions and individual traits (Říha & Prchalová, 2022), impacting predator fitness and lower trophic levels (Shaw, 2020). Recent advances in in situ animal observation methods offer the opportunity to gain a deeper understanding of individual predator movement patterns, their dynamics and underlying factors (Nathan et al., 2022). This progress has improved our ability to identify proximate and ultimate effects at multiple scales, from the individual level to the entire ecosystem (Nathan et al., 2022).

The presence and availability of food are among the most important factors that determine the movement patterns of an individual and impacting home range, activity level or timing (Mitchell & Powell, 2004; Zhdanova & Reebs, 2006). These changes in movement patterns are driven by the energetic cost of foraging (Rennie, Collins, Shuter, Rajotte, & Couture, 2005). Individuals evaluate environmental quality and adjust movements to optimize the balance between feeding gains and costs (Boisclair & Leggett, 2011).

Individual activity levels and home range size are important parameters that are defined based on the interplay among behaviour, physiology, and population density (Haskell, Ritchie, & Olff, 2002). Numerous studies have demonstrated a negative correlation between home range size or activity and prey density (Ferguson, Currit, & Weckerly, 2009; Herfindal, Linnell, Odden, Nilsen, & Andersen, 2005; Loveridge et al., 2009). However, empirical evidence suggests that the relationship is not straightforward and that a hump-shaped relationship may occur when food is scarce (Massei, Genov, Staines, & Gorman, 1997; Scharf, 2016; Spiegel, Harel, Getz, & Nathan, 2013). Such a relationship represents a conservative individual response that minimizes the risk of physiological collapse during starvation.

The availability of food can also influence the timing of activity, regardless of its space use and activity level (Orpwood, Griffiths, & Armstrong, 2006). In fish, studies have shown that when food is scarce, fish adapt their activity patterns and become more active at times when they would normally be inactive to increase encounter rates with prey (Zhdanova & Reebs, 2006). This adaptation helps to compensate for the lack of food and maintain individual growth at a level similar to that observed under food-rich conditions (Orpwood et al., 2006).

In addition to species-specific behavioural patterns, there are significant individual differences in behaviour and movement in response to food availabilityfood availability that are closely related to differences in individual traits of different behavioural types (Campos-Candela, Palmer, Balle, Álvarez, & Alós, 2019; Dingemanse et al., 2007). Theoretical studies propose that different behavioural types show different movement responses when confronted with situations where resource availability is limited. In such instances, the relationship between individual variability in space use and its plasticity should become clearer and space use strategies should be crucial for individual growth and fitness (Campos-Candela et al., 2019). However, there is a lack of empirical evidence linking food availability and inter-individual differences in space use and field observations are important to establish a conclusive link. Furthermore, distinct diurnal patterns, termed chronotypes (Alós, Martorell-Barceló, & Campos-Candela, 2016; Harrison et al., 2015), suggest consistent inter-individual differences in activity timing. Understanding the relationship between food availability and activity timing variation remains limited. It is plausible that in the face of food scarcity, individuals are forced to maintain a consistently higher level of activity in order to increase their foraging efficiency that may alter the differences in activity timing between individuals.

Seasonal changes in movement patterns due to temperature may overshadow the effects of food availability on predator movements. In fish predators, seasonal variations in temperature play a crucial role in various physiological mechanisms, including locomotor capacity and activity levels (Volkoff & Rønnestad, 2020). They can result in reduced activity, especially during the cold season, regardless of food availability conditions. In contrast to this assumption, there are reports suggesting that feeding activity during colder periods may be influenced by body fat reserves (Teletchea et al., 2009). These species may increase their activity during cold periods when their fat reserves are low, but also under conditions of low food availability. However, our knowledge of the behaviour of aquatic predators during the cold season is limited (Říha et al., 2022).

For our study on the effects of food availability on predator’s movement patterns we used the European catfish (*Silurus glanis,* hereafter referred as catfish). This species is one of the largest freshwater predators that is currently attracting increasing scientific interest also in relation to its actual spread across Europe as an invasive species (Encina et al., 2024; Vejřík et al., 2019a). The catfish reaches a considerable body size and exerts a strong influence on ecosystems as an apex predator (Vejřík et al., 2017). Previous studies have mainly described a uniform pattern of movement characterized by long periods of residence and prevailing nocturnal activity (Cucherousset et al., 2018). However, recent studies have revealed contextual variations in movement patterns and individual dietary specialization, shaped by the environment or alteration of activity timing in response to food availability (Bolliet, Aranda, & Boujard, 2001; Slavík & Horký, 2012; Vejřík et al., 2017, 2023). Therefore, it can be expected that food availability in the ecosystem influences the catfish’s intensity of movement, the timing of their activity and space use as well as the resulting individual growth.

The aim of this study is to investigate the ability of catfish as an aquatic top predator to adjust their movements and temporal activity patterns in response to variations in prey density and distribution. Our hypotheses are as follows: i) reduced food availability leads to an increase in activity levels and consequently to expanded spatial use throughout the day during the summer months; ii) the effects of food availability on spatial use and activity will be less pronounced in the colder months; iii) reduced food availability will lead to greater temporal and inter-individual variability in spatial use and activity timing, which will persist in both cold and warm months; iv) increased activity will potentially compensate for lower food availability, leading to similar growth rates in fish prey poor and fish prey-rich environments. To test these hypotheses, we conducted an extensive tracking study of catfish behaviour over a six-month period in three man-made waterbodies with different prey availability and its distribution. Two of these lakes were oligotrophic, with limited food resources, while one was a eutrophic reservoir with abundant food resources.

## Material and Methods

### Study sites

The study was conducted in three artificial waterbodies with comparable size: two oligotrophic lakes, Milada (50° 39′ N, 13° 58′ E; herein referred as OLIGOT1) and Most (50° 32′ N, 13° 38′ E; herein referred as OLIGOT2) and one eutrophic reservoir, Rimov (48°50’N, 14°29’E; Table 1; herein referred as EUT), all located in the Czech Republic. OLIGOT1 and OLIGOT2 are recently created post-mining lakes resulting from aquatic restorations of mining pits (aquatic restoration took place from 2001 to 2010 in OLIGOT1 and from 2008 to 2014 in OLIGOT2). The EUT reservoir was built in 1978 by damming the Malse River. Oligotrophic lakes have a similar fish community composition, with roach (*Rutilus rutilus*) and perch (*Perca fluviatilis*) dominating both lakes, while ruffe (*Gymnocephalus cernua*), tench (*Tinca tinca*), European catfish (*Silurus glanis*), northern pike (*Esox lucius*) and pikeperch (*Sander lucioperca*) being less common. In addition, planktivorous maraena whitefish (*Coregnus maraena*) is present in the open water habitat of OLIGOT2 (Riha et al. 2021). In EUT, roach and bream dominate the fish community, while perch and ruffe are less common (Tesfaye et al., 2022). Previous studies indicated that prey availability for catfish is similar in oligotrophic lakes, with lower prey availability in the open water (Vejřík et al., 2017). On the other hand, in EUT, the littoral and open water prey fish communities are much more numerous (Moraes et al., 2021; Říha et al., 2021;).

**Table 1.** Hydrological parameters of the three study waterbodies. TP stands for total phosphorus concentration in water, SD for Secchi depth and B/P for percentage of water volume in benthic and pelagic habitats.

|  | Area | Volume | Max. depth | Mean depth | TP | SD | B/P |
| --- | --- | --- | --- | --- | --- | --- | --- |
|  | (ha) | (10<sup>6</sup> m<sup>3</sup>) | (m) | (m) | (µg L<sup>-1</sup>) | (m) | (%) |
| OLIGOT1 | 250 | 36 | 25 | 16 | 5 | 5 | 11/89 |
| OLIGOT2 | 311 | 70 | 75 | 22 | 2.5 | 7.7 | 15/85 |
| EUT | 210 | 34 | 45 | 16 | 26.4 | 1.7 | 12/88 |

All tracked catfish in oligotrophic lakes were stocked into these lakes for biomanipulation purposes. Stocking occurred from 2011 to 2013 (2 to 4 years prior to tracking), and catfish from 2012 and 2013 were PIT-tagged prior to stocking. Natural reproduction of catfish was not detected in the lakes before and during the study period. Long-term, regular catfish stocking was conducted in EUT, but catfish also reproduce there naturally (Říha et al., 2009). The stocked catfish were PIT-tagged during stocking after 2012, but the age of the tracked fish could not be determined. Therefore, the origin (natural or stocked) of the tracked catfish could not be distinguished in EUT.

### Fish tagging

A total of 45 individuals (15 in each waterbody) were caught by electrofishing (11 individuals), longlining (33 individuals) and angling (1 individual). Individuals were measured, weighed and sexed prior to tagging (Table 2). Acoustic transmitters - Lotek Wireless Inc, MM-M-11-28-PM, 65 x 28 mm, mass in air of 13 g, including pressure (all waterbodies), motion (oligotrophic lakes) and temperature (EUT) sensors, burst rate 25 s (oligotrophic lakes) or 15 s (EUT), tag weight between 0.03% and 0.6% of fish body weight - were implanted as described in detail in (Říha et al., 2021, 2022). Fish were caught and tagged between 5 and 10 May 2015 in oligotrophic lakes and between 19 and 20 April 2017 in EUT.

**Table 2.**
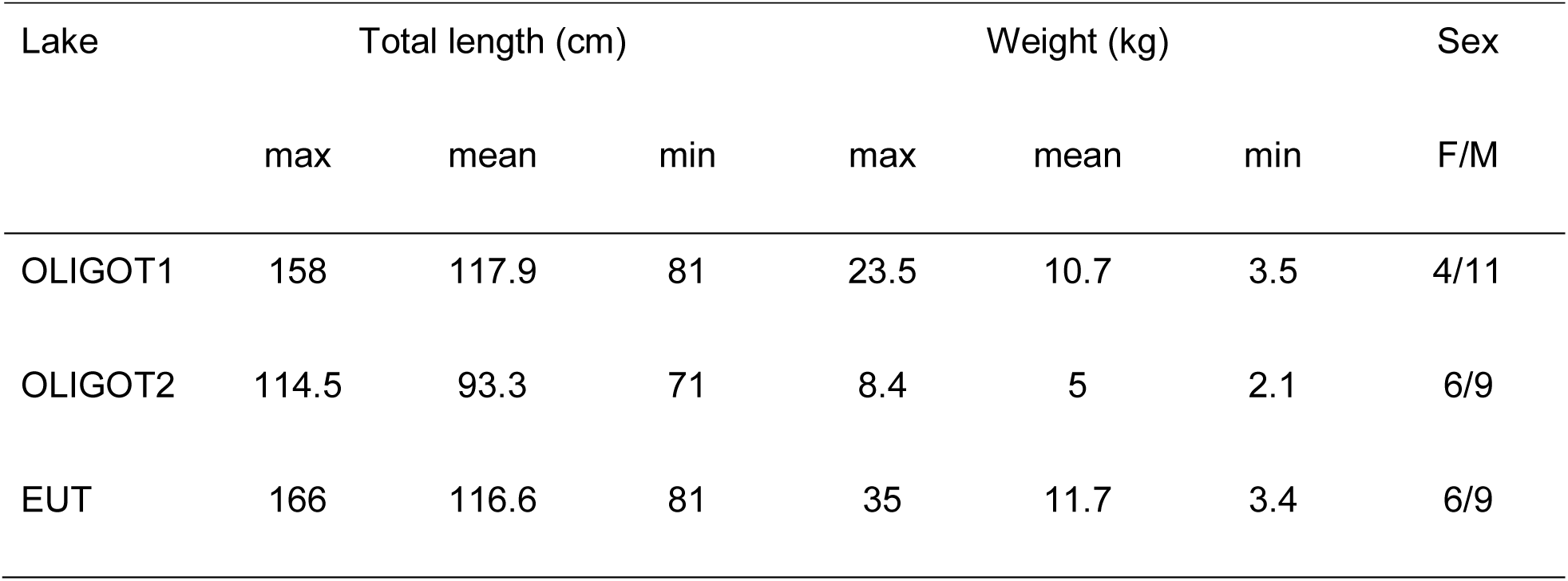
Description of tracked catfish in the three study waterbodies.

| Lake | Total length (cm) |  |  | Weight (kg) |  |  | Sex |
| --- | --- | --- | --- | --- | --- | --- | --- |
|  | max | mean | min | max | mean | min | F/M |
| OLIGOT1 | 158 | 117.9 | 81 | 23.5 | 10.7 | 3.5 | 4/11 |
| OLIGOT2 | 114.5 | 93.3 | 71 | 8.4 | 5 | 2.1 | 6/9 |
| EUT | 166 | 116.6 | 81 | 35 | 11.7 | 3.4 | 6/9 |

### Telemetry system

Three separate MAP positioning systems (Lotek Wireless Inc., Canada) were deployed in all waterbodies to track the tagged fish. The systems consisted of 91 receivers (Lotek Wireless Inc., WHS3250; 44, 47 and 91 receivers in OLIGOT1, OLIGOT2 and EUT, respectively) deployed in arrays (Fig. 1). A detailed description of the design, deployment and accuracy of the arrays can be found for oligotrophic lakes in (Říha et al., 2021) and for EUT in (Říha et al., 2022). For the purpose of this study, tracking records of period June – November were selected (year 2015 in oligotrophic lakes, 2017 in EUT).

**Fig. 1.**
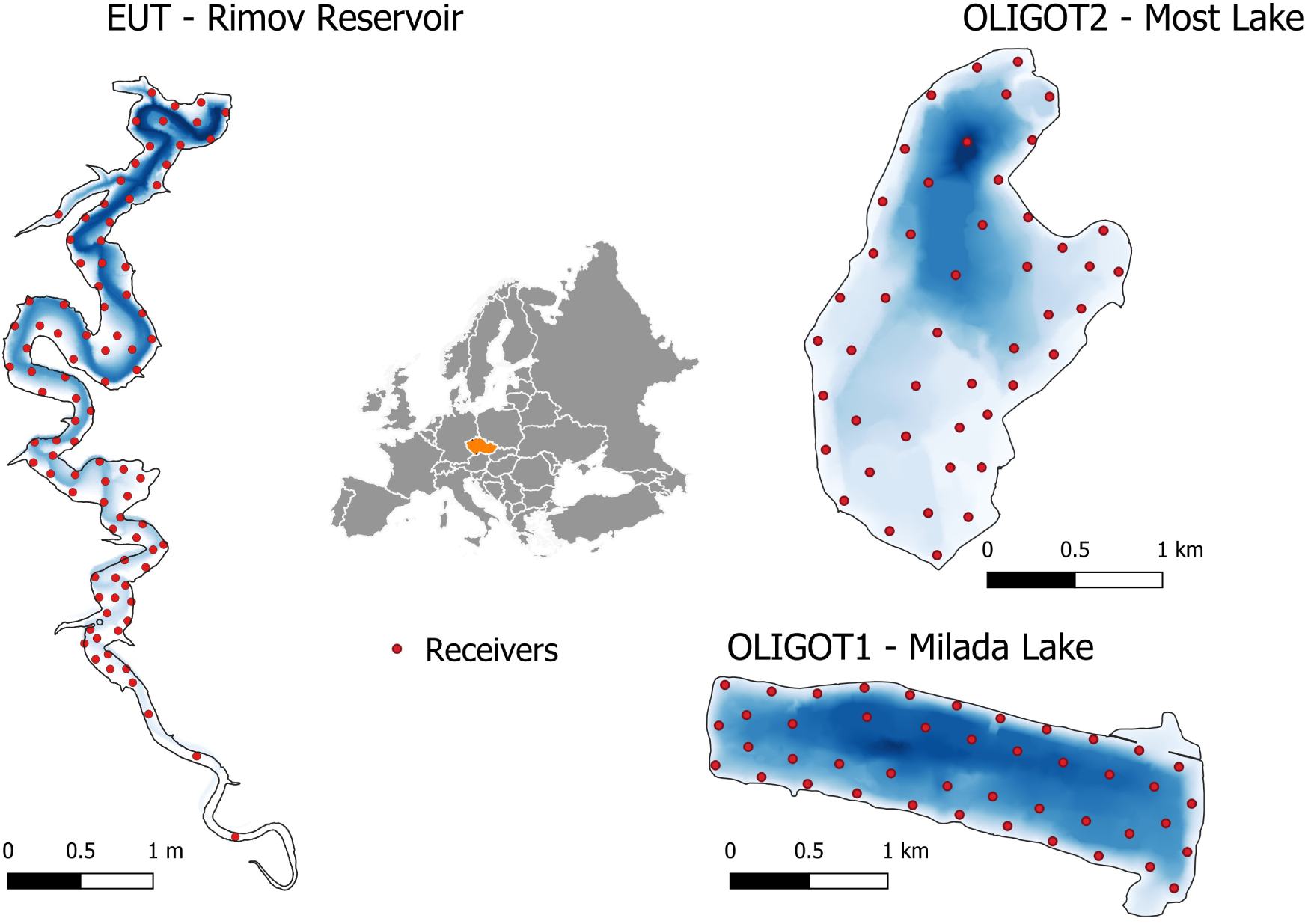
Locations and bathymetry (blue color) of the investigated waterbodies and positions of the telemetry systems. The orange color indicates the position of the Czech Republic where the study was conducted. The dots represent the positions of the individual telemetry receivers.

### Catfish growth, population size and diet sampling

Catfish populations were long-term monitored in all three waterbodies, and all caught catfish were PIT-tagged and released back into the waterbody (see (Vejřík et al., 2017, 2023) for more details). Recaptures of PIT-tagged catfish were used to calculate individual catfish growth. Growth was calculated as the difference between the length at the time of PIT tagging and the last recapture scaled with time (length gain/year). If the time interval between capture and recapture was less than 1 year, growth was not calculated.

The stomach contents of the catfish were removed by hand through the open mouth and gullet. The stomach contents were then identified, measured and weighed or fixed with 70% ethanol for laboratory identification using diagnostic elements such as fish bones (for more details see (Vejřík et al., 2017).

### Fish community sampling

Catfish fish prey availability and distribution were investigated using stratified fish community sampling by gillnet and hydroacoustic surveys. Gillnet surveys were conducted in benthic habitats (2 sites in oligotrophic lakes and 4 sites in EUT) and open water habitats (1 site in oligotrophic lakes and 4 sites in EUT), using 30 m long European standard gillnets with multiple meshes set overnight (more details in (Říha et al., 2021; Tesfaye et al., 2022)). For this study, only catches of fish older than fingerlings were considered. Gillnet surveys were conducted in oligotrophic lakes in September 2015 and in EUT in August 2017.

Density as catch per unit effort was measured as the number of fish caught per 1000 m2 gillnet area per night (NPUE). NPUE was then weighted by water volume in open water and benthic habitats (Table 1) to give a more realistic approximation of the differences between distribution of fish densities between these two habitats (Tesfaye et al., 2022).

The Simrad EK 60 echosounder was used for the hydroacoustic evaluation of the fish stocks. Two Simrad ES 120-7C transducers, operating at a frequency of 120 kHz, were set to cover the entire water column simultaneously (horizontally and vertically) during day and night (Draštík et al., 2009; Říha et al., 2021). In the Sonar5-Pro software (CageEye AS, Oslo, Norway), echointegration was used to estimate the density of individual fish while shoals were detected manually. Volume backscattering thresholds were set to –62 dB (40logR) and target strength thresholds were set to –56 dB, corresponding to a fish length of 4 cm. The fish tracks were used as the source for calculating the mean target strength.

### Data processing

Locations of individual fish were calculated using proprietary post-processing software UMAP v.1.4.3 (Lotek Wireless Inc., Newmarket, Ontario, Canada). Filtering and further postprocessing of fish locations are described in detail in (Říha et al., 2021).

The extent of horizontal area use was calculated using a 95% kernel utilization distribution (hereafter referred to as KUD). Daytime and nighttime were defined as the time before and after sunset and sunrise, respectively. The exact time of sunset and sunrise on each day was calculated using the R package “maptools” (Bivand & Lewin-Koh, 2015) and UD was calculated using the R package “adehabitatHR” (Calenge & Fortmann-Roe, 2019). Further details of UD calculation are given in (Říha et al., 2021). The activity of fish was calculated as horizontal swimming speed (expressed in meters per second) between two consecutive positions.

To test the importance of an open water habitat for catfish in all three waterbodies, the proportion of time spent in open water was calculated (TOW). Each mean position was assigned to be either in the benthic (distance < 5 m from the bottom) or in the open water habitat (≥ 5 m from the bottom) and the ratio between the number of open water and benthic locations was calculated.

### Statistical analysis

To investigate catfish space use and activity in the three waterbodies, we created a Generalized Additive Mixed model (GAM). This model incorporated the daily time series (time) spanning a six-month period as a smoothing component, with waterbody identity coded as a parametric factor. Additionally, to address residual autocorrelation, we integrated a serial autocorrelation into the model, ensuring independence from immediately adjacent observations. We defined both monthly (*month*, starting in June) and time of the day (*tod*, range 0-24 hours) variables to account for seasonality during the study period. Waterbody identity was also included as a by-group variable to fit different temporal trends by each waterbody. Together with catfish body length and sex, both seasonal components, *tod* and *month,* were fitted with thin plate and cubic spline functions, respectively. The suitability of including an interaction between the seasonal components (*tod*-by-*month*) was tested with different smoothing and tensor product terms. In separate models using a subset of the data from each waterbody, we accounted for differences in fish trajectories (slopes over time) by including random smooth factors for each fish identity tag (fishID), which allowed us to estimate the temporal change as a function of variability within each waterbody (see Supplementary material for more details). The models were fitted using the *bam() f*unction from the R package “mgcv” (Wood, 2021) using a restricted maximum likelihood method, adopting a Gaussian error distribution and identity link function. Model comparisons were conducted via likelihood ratio tests, where minimal variance differences favoured the simplest model. We tested the consistency of the behaviour using the GAM models as in (Jarić et al., 2022).

For analyzing of open-water habitat use (TOW), we employed Generalised Additive Models for Location, Scale and Shape (GAMLSS) (Stasinopoulos & Rigby, 2007) fitted using the R package “gamlss” (Rigby & Stasinopoulos, 2005) with a one-inflated beta distribution. TOW encompasses values in the interval [0,1], reflecting varying levels of open water habitat use (i.e., TOWP), where a value of 1 indicates maximal open water utilization (i.e, TOWt). Models were fitted separately for each waterbody, comprising three distribution parameters: the mean *μ,* variance (scale) *σ*l, and *v* denoting the point probability at 1 (detailed parameterization of the GAMLSS models is available in the Supplementary material and Říha et al., 2021). In the TOW model, we incorporated the interaction between time, fitted with a cubic smoothing spline, and diel period to account for temporal variability across waters. Additionally, body length was included with a penalized P-spline to address potential size effects. -Model selection was guided by the Schwartz information criterion (SBC) values, to select the smooth terms and the composition of each distribution parameter (see Supplementary material for more details). A random smoothing function accounted for interindividual variability, and an autocorrelation structure accounted for residual correlation.

## RESULTS

### Daily horizontal space use

Daily horizontal space use (HSU) was highest in OLIGOT1 (mean ± SE: 22.47 ± 2.59 ha/day, t = 8.67, P < 0.001), followed by OLIGOT2 (16.97 ± 2.62 ha/day, t = 6.48, P < 0.001), and the lowest in EUT (14.64 ± 2.61 ha/day, t = 5.61, P < 0.001; Fig. 2a). In OLIGOT1, HSU was higher than in EUT and OLIGOT2 throughout the whole period, with the exception of the latter lake in July, when HSU in OLIGOT2 was comparable to or slightly higher than in OLIGOT1 (Table 3, Fig. 2b).

**Fig. 2.**
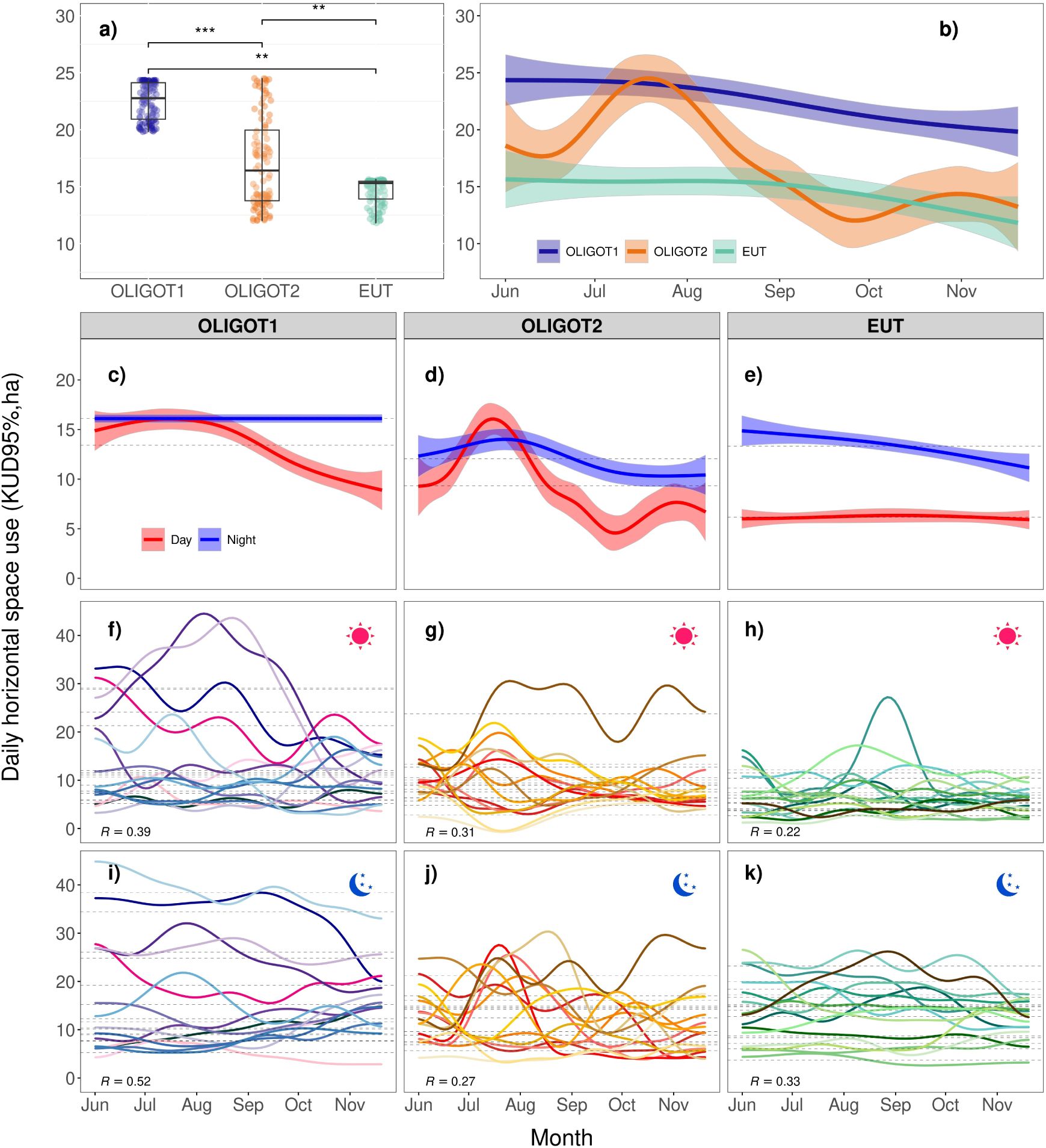
a) Comparison of mean catfish daily horizontal space use between the waterbodies. The asterisks indicate the significance of the differences (* p < 0.05, ** < 0.01, *** < 0.001) ; b) seasonal and c-e) diel development of the mean daily horizontal space use estimated by the GAM models; individual trajectories of horizontal space use during the day (f-h) and at night (i-k) estimated by the GAM models. R, adjusted repeatability index calculated from the final fit model. The dashed lines (i-k) show the average individual mean daily horizontal space use.

**Table 3.** Differences in mean activity and spatial use of fish in lakes OLIGOT1, OLIGOT2 and EUT from June to December, modelled by generalised additive mixed models (GAMs). The adjusted R2 and the scale parameter (ϕ) were estimated directly from the model, the latter being associated with the residual variance. Signif. codes: * p<0.05; ** p<0.01; *** p<0.001.

|  | HSU |  | Mean activity |  |
| --- | --- | --- | --- | --- |
| *A. Parametric coefficients* | Estimate (S.E) | t-value | Estimate (S.E.) | t-value |
| (Intercept) | 22.46 (2.59) | 8.67*** | 0.08 (0.005) | 14.298*** |
| OLIGOT2 | -5.51 (3.68) | -1.50 | -0.01 (0.01) | -0.789 |
| EUT | -7.82 (3.68) | -2.13* | -0.001 (0.01) | -0.123 |
| *B. smooth terms* | Edf (Ref. df) | F-value | Edf (Ref. df) | F-value |
| s(Time) × OLIGOT1 | 1.99 (9.00) | 2.03*** | 6.93 (9.00) | 128.40*** |
| s(Time) × OLIGOT2 | 6.05 (9.00) | 11.70*** | 7.08 (9.00) | 66.60*** |
| s(Time) × EUT | 1.77 (9.00) | 2.00* | 7.27 (9.00) | 42.65*** |
| s(bl) |  |  | 1.95 (9.00) | 3863*** |
| te(month, tod):OLIGOT1 |  |  | 87.24 (137.00) | 10.15*** |
| te(month, tod):OLIGOT2 |  |  | 45.82 (137.00) | 3.83*** |
| te(month, tod):EUT |  |  | 113.60 (137.00) | 34.30*** |
| s(fishID) | 40.43 (42.00) | 33.92*** | 38.093 (41.00) | 36.37*** |
| Adjusted R<sup>2</sup> | 0.42 |  | 0.22 |  |

Overall, the nocturnal HSU in the three waterbodies was higher than the daytime HSU (Table 4). During the night, HSU was highest in OLIGOT1 and lowest in OLIGOT2 (P < 0.001; Fig. 2c-e). The difference between day and night was particularly pronounced in EUT (P < 0.001) and significantly lower in the two oligotrophic lakes (both P < 0.001, Table 4). In EUT catfish used significantly less horizontal space during the day than in oligotrophic lakes (P < 0.001, Table 4, Fig. 2c-e). Individual and inter-individual variation was lower in EUT than in the oligotrophic lakes (Table 5; Fig. 2f-k). Comparisons between day and night reflected this pattern, with EUT showing the lowest repeatability (Table 5; Fig. 2f-k).

**Table 4.**
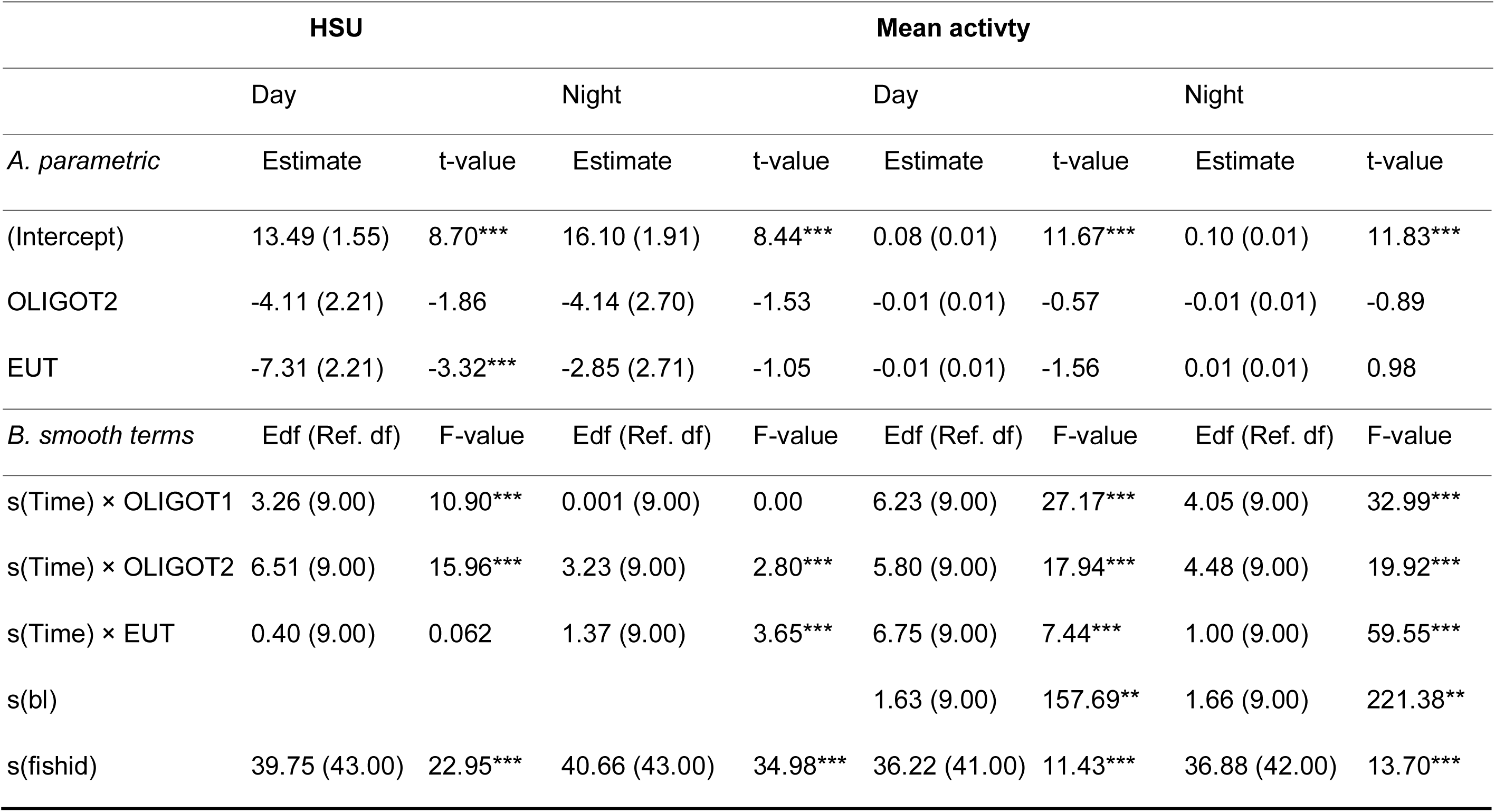
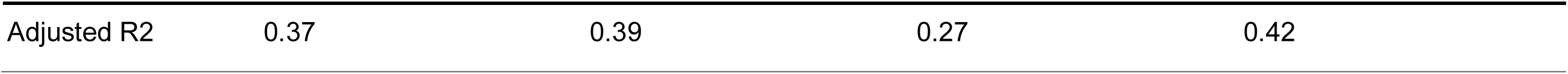
Model differences in HSU and mean activity between day and night. Signif. codes: * p<0.05; ** p<0.01; *** p<0.001.

**Table 5.**
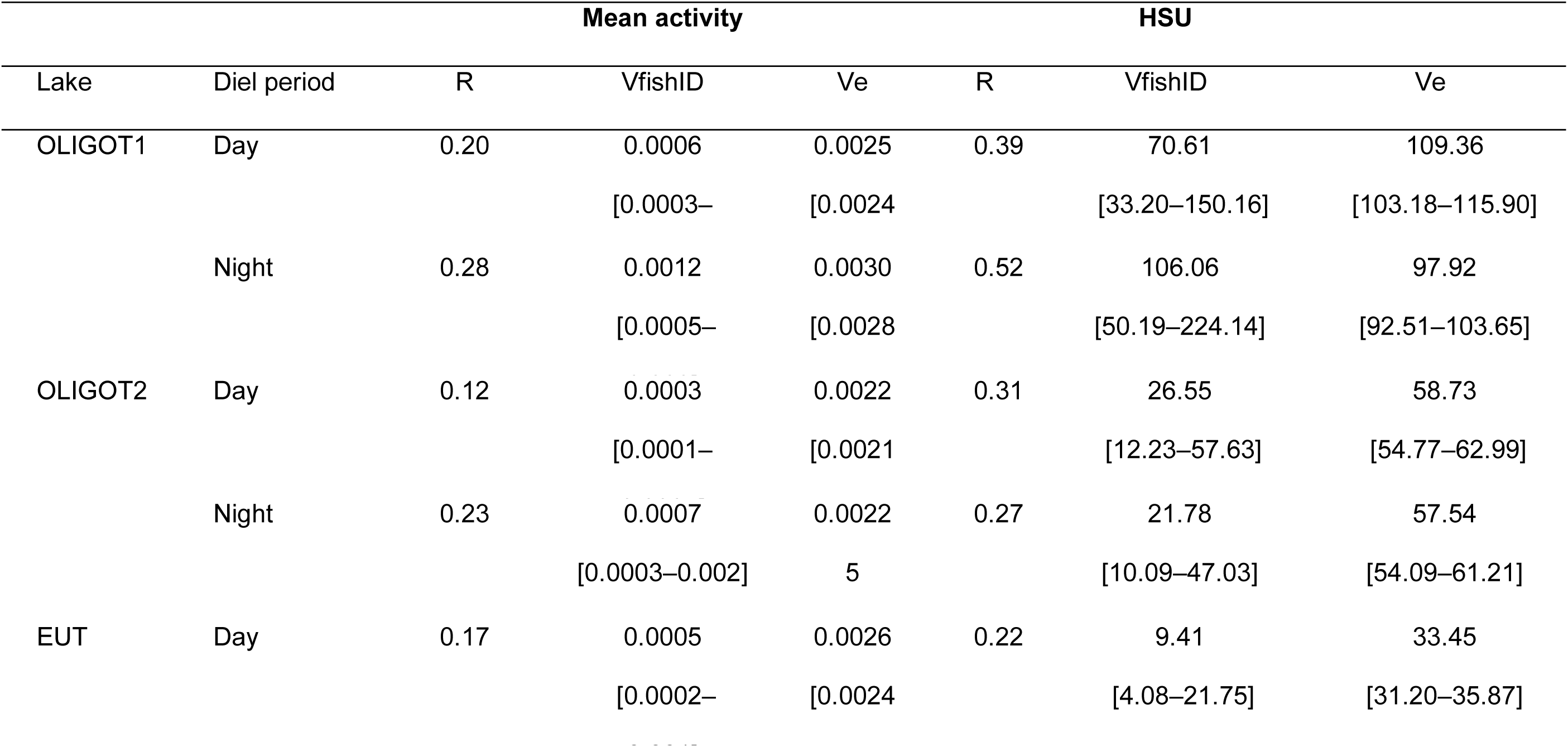
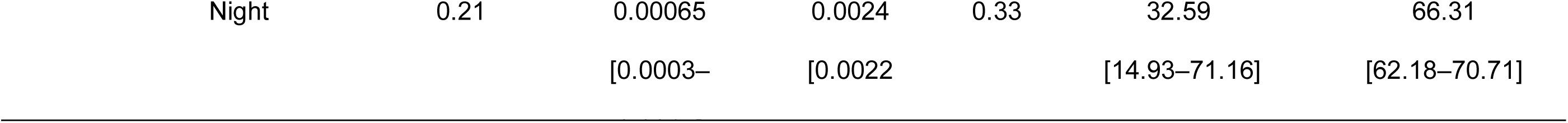
Repeatability of mean activity and daily horizontal space utilization of fish by lake and time of day. Data show only the adjusted (*Rslope*) R-values calculated from the final fit model, with 95% confidence intervals calculated using GAMs of increasing complexity, starting with random intercepts and including fixed effects and random slopes. The variance components are also shown for the selected model. (see SM for more details on the calculations of the variance and R indices).

### Activity

Catfish activity changed non-linearly during the study period, and overall activity was similar in all waterbodies (Table 3), with a peak from June to August and a subsequent decline (Fig. 3a). The average swimming activity was very similar in OLIGOT1 and EUT and lowest in OLIGOT2 without significant difference (Fig. 3a). Diurnal activity varied considerably between waterbodies (Table 4), especially from June to September, mirroring the pattern in HSU (Fig. 3 b-g). In EUT, catfish were less active during the day and doubled their activity at night (Table 4, P < 0.001), while no such large differences between day and night were observed in the two oligotrophic lakes (Table 4, Fig. 3 b-g).

**Fig. 3.**
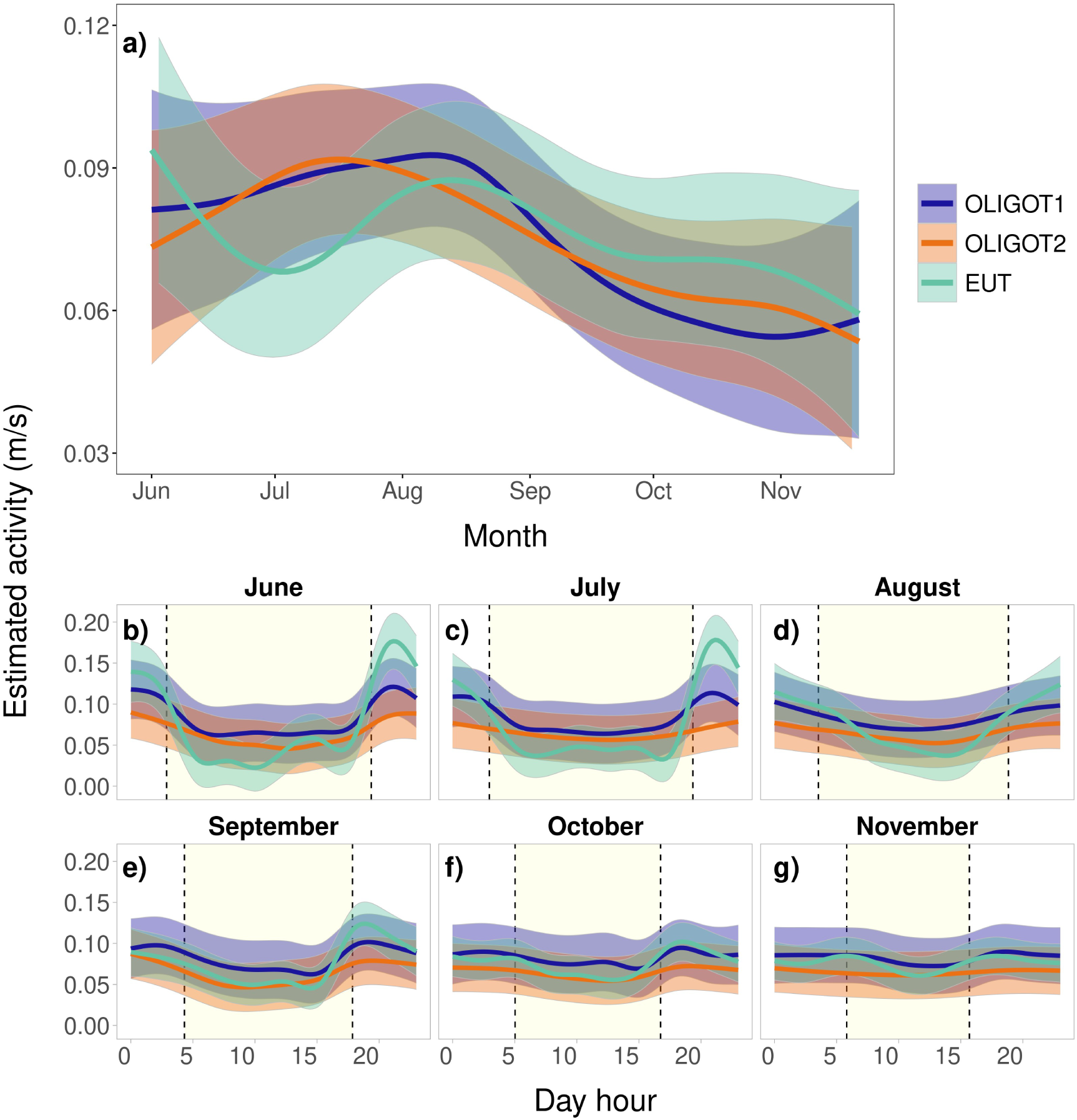
a) Mean catfish swimming activity during tracking period in the waterbodies studied; b - g) effect of time of day on swimming activity in each month and waterbody, with the mean time of sunset and sunrise (dashed lines) and the time of daylight in yellow.

Average activity in the three waterbodies varied to a similar extent between and within individuals, both during the day and at night, with individual variability generally greater than inter-individual differences. Only in OLIGOT1 the individual differences between day and night were significant, being twice as high at night as during the day (Table 5; Fig. S1).

### Habitat use

In general, catfish used the open water only partially (waterbody(μ); p(0 > TOW < 1with a proportion roughly comparable between the two oligotrophic lakes (TOWP = 0.31 and 0.34, respectively) and higher in EUT (TOWP = 0.48; Table 6, Fig. S3). The probability of catfish fully utilizing the open water habitat (waterbody(ν); p(TOW = 1)) was highest in OLIGOT2 (TOWt = 0.10), followed by EUT (TOWt = 0.04) and OLIGOT1 (TOWt = 0.03). In OLIGOT1, there were no significant overall differences in the TOWP rates between day and night (diel period(μ), day = 0.32; night = 0.36, *P* = 0.15), although the temporal trends tended to diverge in opposite directions (Table 6; Fig. 4). In OLIGOT2, the TOWP proportion was higher at night than during the day (diel period(μ), day = 0.27; night = 0.34, *P* < 0.01) and temporal patterns did not differ significantly. In EUT, TOWP was higher at night than during the day (diel period(μ), day = 0.44; night = 0.52, *P <* 0.01), and these differences persisted over studied period, with a tendency to increase during the day and decrease at night (Table 6; Fig. 4, S3).

**Fig. 4:**
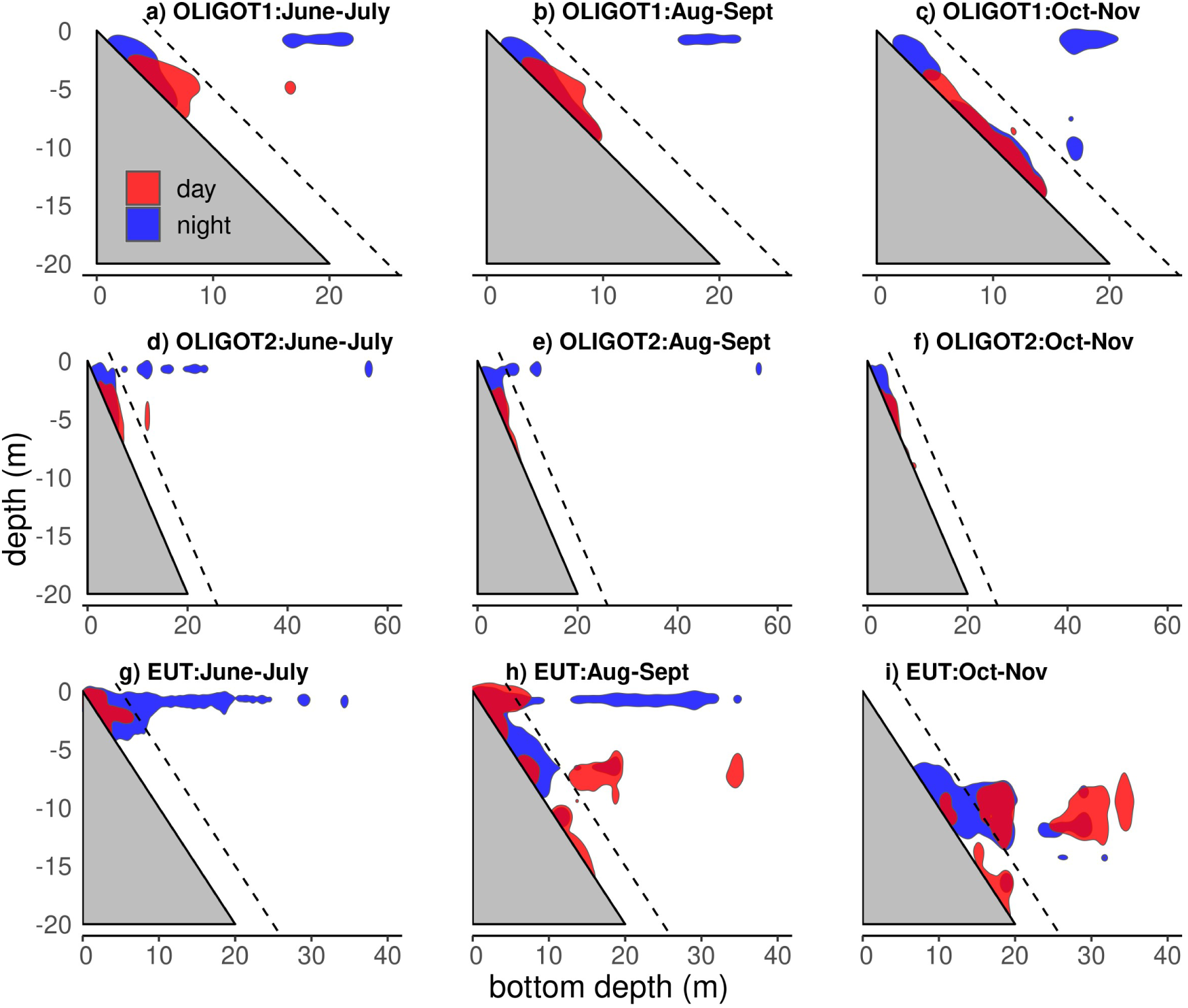
Utilization of depth by catfish in relation to bottom depth in each waterbody and at two- month intervals (polygons representing a 50% core utilization distribution). The dashed line shows the threshold of 5 m used for the calculation of open water utilization (TOW parameter).

**Table 6.**
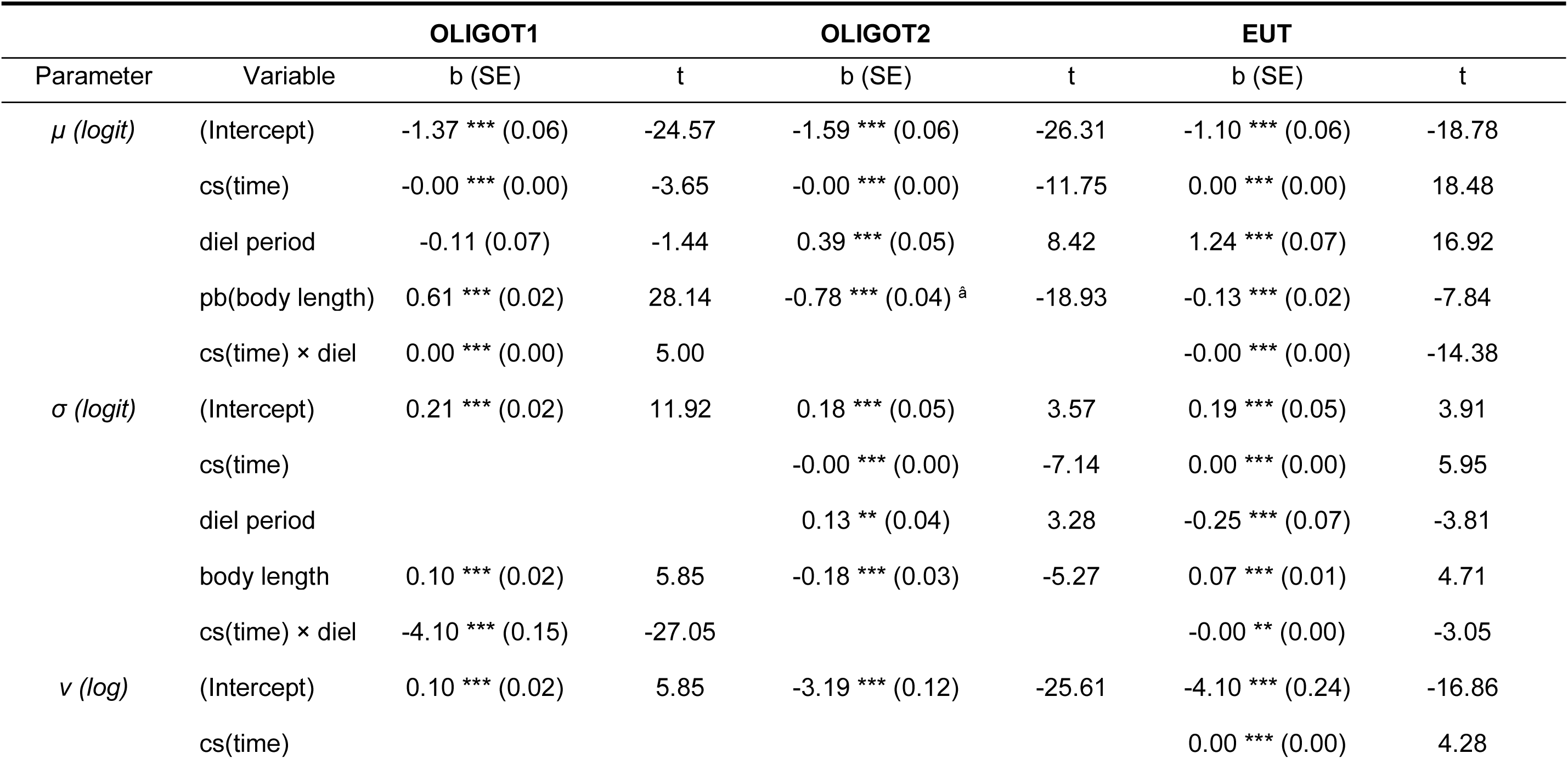
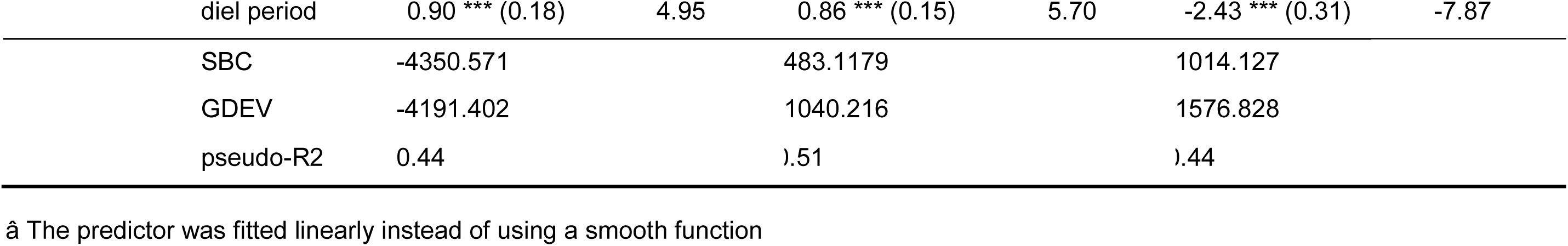
GAMLSS analysis of pelagic habitat use in three lakes as a function of the proportion of time spent in open water (TOW). b, variable estimate (z-score); SE, standard error; t, t-statistic; GDEV, global deviance; SBC, Schwartz Bayesian Criterion; pseudo- R^2^, generalized pseudo-R-squared. Signif. codes: * p<0.05; ** p<0.01; *** p<0.001..

### Growth and diet of catfish

Individual growth varied significantly between the waterbodies (ANOVA, p < 0.001), with EUT showing the highest growth (Fig. 5a). In all waterbodies, fish were the main food of the catfish, with the catfish in EUT concentrating almost exclusively on fish. In the oligotrophic lakes, the catfish diet was more diverse and included to a greater extent crayfish in OLIGOT1 and birds in OLIGOT2 (Fig. 5).

**Fig. 5:**
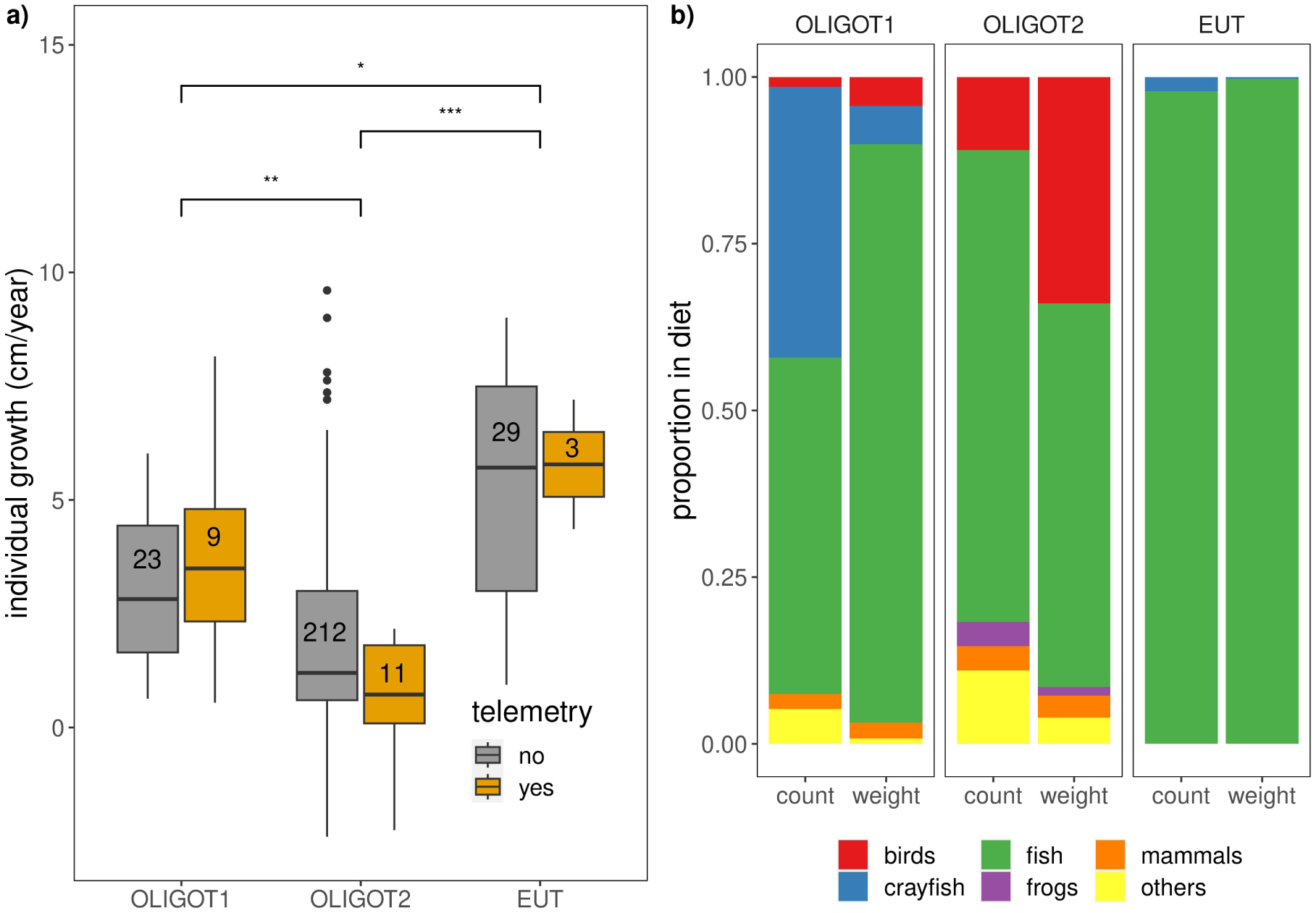
a) Individual body growth defined as increment of body length (cm) per year, yellow boxes show the growth of fish used for the telemetry study, grey boxes show individuals caught during catfish surveys in 2013-2017 in oligotrophic lakes, and 2016-2019 in EUT. The numbers in the boxes indicate the number of individuals recaptured, (* p < 0.05, ** < 0.01, *** < 0.001); b) composition of catfish prey from stomach content analysis.

### Fish prey availability and distribution

In EUT, the fish gillnet NPUE was considerably higher compared to oligotrophic lakes (Fig. 6). The main difference was observed in open water habitats, with 52 to 88 times higher NPUE in EUT. Hydroacoustic estimates yielded similar results with higher open water fish density in the reservoir in both diel periods (Fig. 6). OLIGOT2 had a slightly higher both gillnet and hydroacoustic abundance of fish than OLIGOT1 in both habitats and diel periods (Fig. 6).

**Fig. 6:**
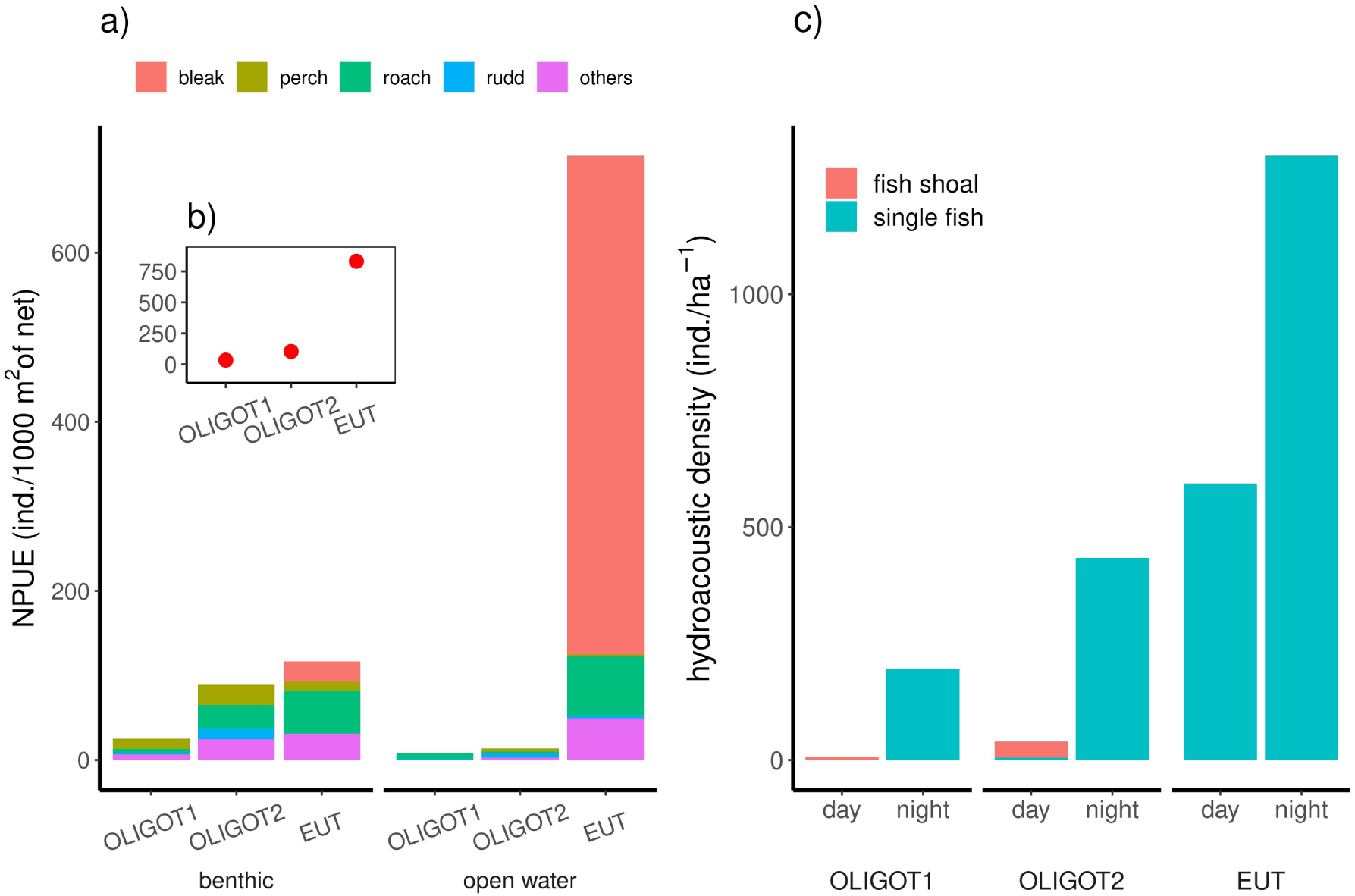
Prey fish abundance in different habitats and times of day: a) weighted (by volume) gillnet NPUE of prey fish in the open water and benthic habitats; b) NPUE estimate for the whole water body; c) estimate of mean hydroacoustic fish density in the waterbody in both times of day

## Discussion

Our study shows that catfish exhibit different spatial movement behaviour in oligotrophic lakes with low fish prey abundance than in a eutrophic reservoir rich in fish prey. According to our predictions, the horizontal space use and diurnal activity patterns differed between waterbodies. In oligotrophic lakes, catfish increased their diurnal activity and space use and decreased their nocturnal activity (hypothesis 1). The differences in the distribution of fish prey between waterbodies also led to a differentiation in catfish habitat utilization, with the open water zone in EUT being used more intensively. These behavioural differences decreased in the colder months (hypothesis 2). The variability of space utilization among and within individuals was higher in the oligotrophic lakes than in EUT, but seasonal development of these activity parameters was similar in all waterbodies (hypothesis 3). The change in diurnal behaviour in the oligotrophic lakes did not fully compensate for the lower food availability, resulting in lower catfish growth in these lakes (hypothesis 4). In addition, catfish prey was more diverse in the lakes with lower food availability.

### Space use and activity

Space use and activity differed mainly during the day in summer between poor and rich fish prey systems, when catfish in oligotrophic lakes showed comparable activity to that at night and used a similar horizontal space. In EUT on the other hand, the catfish were mostly inactive during the day and significantly increased their space utilization and activity at night.

The observed differences in space use and activity patterns among waterbodies could represent an adaptive strategy to optimise resource use in response to differences in prey availability. Catfish are nocturnal species (Boujard, 1995; Pohlmann, Grasso, & Breithaupt, 2001) but able to adapt their activity to the time of higher prey availability as documented in laboratory conditions (Bolliet et al., 2001; Boujard, 1995). Prey abundance showed large differences in our study, with the density of fish prey in the eutrophic reservoir being between 8 and 25 times higher (in terms of gillnet abundance) than in the two oligotrophic lakes. Alteration of catfish diurnal activity patterns in oligotrophic lakes is therefore most likely due to the overall lower prey abundance, which wasinsufficient to satisfy the catfish’s energy requirements and forced the catfish to forage during the day as well.

Steady low-intensity movement in oligotrophic lakes during the day could help catfish to explore new areas and maximize the likelihood of finding resources and food. Moreover, daytime catfish foraging might be important for hunting non-piscivorous prey such as birds, which are an important part of diet in the two lakes. Previous research showed effective diurnal catfish bird hunting (Cucherousset et al., 2012). The bird species found in the stomachs of catfish in our study were coots (*Fulica atra*), cormorants (*Phalacrocorax carbo*) and great crested grebes (*Podiceps cristatus*), species that are only available during the day as their nocturnal refuge is not accessible to catfish (Johansen, Barrett, & Pedersen, 2001; Piersma, Lindeboom, & van Eerden, 1988; Salathé & Boy, 1987).

### Seasonal development of activity and space use

Our results showed similar seasonal development of space use and activity. Previous research has indicated that catfish activity peaks within a temperature range of 25-27°C, sharply declining below 10°C (Britton, Pegg, Sedgwick, & Page, 2007; Copp et al., 2009). Our data corroborate these findings showing that activity and space use peak in summer and subsequently level off by November in all waterbodies, with activity decreasing to 60-80% compared to summer, while the decrease in space use was comparatively lower. Oligotrophic lakes showed a greater reduction in space use during the day, while OLIGOT2 and EUT showed a minimal reduction or, in the case of OLIGOT1, no reduction during the night. These results indicate different movement patterns, with both activity and space use decreasing in the oligotrophic lakes during the day, while night activity was less intense but covered a similar area in all waterbodies. While such patterns suggest territorial behaviour during the night, which is consistent with the notion that catfish are territorial (Carol, Zamora, & García-Berthou, 2007; Daněk et al., 2016), our results showed dynamic seasonal spatial preferences. These preferences were reflected in changes in the open water and depth uses, which did not suggest that they maintained the same territory in the long term. In addition, the marked reduction in diurnal activity and space use in oligotrophic lakes could indicate less profitable foraging opportunities during the day or the absence of certain prey such as migratory birds, amphibians or small mammals. In order to gain a more comprehensive understanding of the seasonal development of catfish behaviour, further detailed studies are needed to investigate movement patterns, habitat use and foraging behaviour.

### Use of habitat

The occurrence of catfish in the open water habitat was influenced by both the time of day and the characteristics of the waterbody. In oligotrophic lakes, the open water was less important for catfish in general, as they preferred shallow benthic habitats both during the day and at night. These preferences are strongly consistent with the distribution patterns of the prey fish. In oligotrophic lakes, the density of prey fish in the open water was conspicuously low during the day, while the density of open water fish increased at night. Despite this nocturnal shift, the gillnet results indicated that fish densities remained higher in the benthic habitats and the probability of catfish using the open water increased only slightly. In the eutrophic reservoir, catfish showed a preference for benthic, shallow habitats during the summer days. This preference was probably less influenced by prey distribution, as catfish were less active during the day and numerous studies have found that catfish rest near underwater structures during the day (Copp et al., 2009). During the night, however, the probability of using the open water increased significantly, with similar utilization rates observed as in the benthic habitats. While the overall density of prey fish was much higher in the open water, the predominant species in the stomachs of catfish, such as roach, perch and bream, had similar densities in both habitats or were even more abundant in the benthic habitats (perch; our results, also (Muška et al., 2018; Říha et al., 2015).

In autumn, the catfish in the eutrophic reservoir moved to deeper water layers and also displayed increased preference for the open water during the day. Such behaviour, which was not observed in oligotrophic lakes, can be attributed to differences in seasonal prey distribution or other environmental factors. Future studies should further investigate the seasonal dynamics of catfish habitat use.

Interestingly, our study showed that catfish were more likely to exclusively use the open water in OLIGOT2 compared to other waterbodies. The exclusive use of open water was individually dependent (see Fig. S3), as only a few individuals had a higher probability of using this habitat. These results are consistent with the general trend towards diversification of foraging under limited resources (Campos-Candela et al., 2019) and highlight greater inter-individual behavioural differences in oligotrophic lakes (see below). Overall, these results emphasize the contextual dependence of prey distribution on predator forage behaviour and support previous research indicating the ability of predators to dynamically change their habitats according to prey availability (Brodersen, Howeth, & Post, 2015; Cruz-Font, Shuter, Blanchfield, Minns, & Rennie, 2019).

### Inter- and intra-individual variability

Although our results suggest that oligotrophic lakes exhibited higher individual variability in space use in comparison to the eutrophic reservoir, OLIGOT1 showed greater individual diversification until early November, while in OLIGOT2 this was mainly the case during the summer. In both lakes and during both periods of the day, individuals divided their space utilization strategies into mobile and sedentary patterns and maintained them consistently. In the eutrophic reservoir, the space use of individuals depended on the time of day. Individuals pursued a very consistent strategy of low space utilization and movement during the day and used the space differently at night by adopting a sedentary or mobile foraging strategy to a similar extent as in oligotrophic lakes.

Theoretical studies suggest that in a prey-poor environment, metabolically fast, mobile animals need to increase their space use, to meet higher energy requirements compared to slow, resident animals (Campos-Candela et al., 2019), as such inter-individual differences become more apparent. Our results are largely consistent with this assumption. However, if only food availability was responsible for the variability of individual space use, we would expect similar variability in space use in the two oligotrophic lakes. The study of (Vejřík et al., 2023) has shown that oligotrophic lakes exhibit a high degree of individual diet specialization, in contrast to the situation in the eutrophic lake. An important part of the catfish diet were birds in OLIGOT2 and crayfish and birds in OLIGOT1. The feeding strategies and habitat should differ greatly for the individuals that specialize on this prey, and this suggests that other mechanisms, such as prey type or distribution, may contribute to the individual differences in these lakes.

### Diet of catfish and food availability

Diet analysis revealed a clear difference in the feeding habits of catfish between the reservoir and the oligotrophic lakes. In the reservoir, catfish favoured almost exclusively fish-type prey, whereas in the oligotrophic lakes they had a broader diet including largely birds (OLIGOT2) or crayfish and birds (OLIGOT1). Although catfish are primarily piscivores (Copp et al., 2009), numerous studies have highlighted their ability to adapt to different food sources and exhibit generalist feeding behaviour (Cucherousset et al., 2018). These results suggest that catfish opportunistically target abundant prey to fulfill their energy requirements. Obviously, the versatility of foraging behaviour, coupled with the adaptability of behaviour, allows catfish to use a wide range of prey and diversify their foraging strategies. This foraging flexibility means that catfish in oligotrophic lakes forage throughout the day and adapt their foraging habits to take advantage of dynamic changes in food supply. In contrast, catfish in the eutrophic reservoir appear to conserve energy during the day and actively hunt for prey, especially forage fish, at night.

### Growth

Our results showed that diet and behavioural flexibility of catfish in oligotrophic lakes did not fully compensate for reduced food availability, leading to reduced catfish growth. There are several possible explanations for this pattern, with several factors possibly interacting to reduce growth rates. The lower prey density observed in oligotrophic lakes could reduce the encounter rate of catfish with their prey and thus their hunting success, which could otherwise lead to increased foraging activity. In addition, daytime hunting could be less effective overall. Catfish are primarily nocturnal feeders relying heavily on senses other than sight, such as lateral line detection and prey wave detection, as their vision is limited (Pohlmann et al., 2001). Consequently, fish prey in oligotrophic lakes with high transparency may exhibit increased avoidance behaviour, which further impairs the hunting success of catfish. Furthermore, changes in diel activity patterns can mean additional energy expenditure for catfish. (Slavík & Horký, 2012) documented that catfish with increased daytime activity expended relatively more energy than catfish that adhered to a more typical diel activity pattern. In addition, catfish grew slower in OLIGOT2 than in OLIGOT1, indicating poorer conditions in this lake. Like birds in OLIGOT2, crayfish were also an important part of thecatfish’s diet in OLIGOT1. Crayfish are a smaller and (according to our observations) more abundant prey than fish and birds and represent a readily available prey source for the catfish in the lake. Their higher utilization could compensate to some extent for the lack of fish prey and the increased growth of catfish. These results suggest that the behavioural adaptations of catfish to prey type and density may help to offset the effects of limited prey resources.

### Caveats

We suggest that the significant differences in fish prey abundance are the most plausible explanation for the observed differences in catfish behaviour and growth in the different waterbodies we studied. However, it is important to point out that our study was an uncontrolled experiment conducted without replication in the prey rich reservoir environment. Therefore, we were not able to establish a direct causal relationship between food availability and the observed behavioural differences between catfish. In addition, our study sites differed in various environmental parameters such as structural complexity, water transparency, maximum depth, prey species composition and oxygen availability. These differences could influence catfish behaviour in ways other than just prey availability. In addition, catfish were released into all waterbodies and previous research has shown increased space utilization by catfish introduced to the new environment (Monk et al., 2020). However, the study by (Monk et al., 2020) indicates a persistent increased activity for months after stocking, while the catfish in our study had already been living in the waterbodies for at least 2 years after stocking and thus had sufficient time to adapt to the new environment.

### Catfish impact

The results underline the remarkable behavioural and feeding flexibility of catfish, which enables them to cope effectively with different environmental conditions and adapt quickly to local conditions. This flexibility encompasses not only the interactions between diet and feeding behaviour but also the plasticity of individual behavioural traits. The optimisation of catfish activity and space use strategies observed in our study highlights the evolutionary need to maximise foraging efficiency and overall fitness in different habitats, giving catfish a significant competitive advantage. This adaptability also probably contributes to the rapid spread of catfish populations in new areas, which has a significant impact on the invaded ecosystems (Vagnon, Cattanéo, Goulon, Guillard, & Frossard, 2022; Vejřík et al., 2019b). This phenomenon is particularly evident in the context of global warming, where rising temperatures, which were once a limiting factor for catfish dispersal, are now facilitating their spread into previously inaccessible regions (Britton, Cucherousset, Davies, Godard, & Copp, 2010).

Our results also suggest significant effects on the risk for prey to be preyed upon by catfish, which depend on the environmental context. The spatio-temporal differences in catfish activity and habitat utilization between the oligotrophic lakes and the eutrophic reservoir led to different landscapes of fear within the ecosystems. These differences can have profound effects on both direct and indirect predator-prey dynamics (Gallagher, Creel, Wilson, & Cooke, 2017), potentially impacting not only prey populations but also other trophic levels (Laskowski, Nuñez, Hilt, Gessner, & Mehner, 2022); Lichtenstein et al. 2023). To date, there is a lack of studies investigating the effects of the environmentally dependent behaviour of apex predators on ecosystems. Therefore, further research is needed to better understand the nuanced interplay between predator behaviour and ecosystem dynamics, especially in the context of changing environmental conditions. This knowledge is crucial for the development of more comprehensive management and conservation strategies aimed at maintaining ecosystem integrity and biodiversity in the face of ongoing environmental change.

## Conclusion

Our study highlights the remarkable behavioural flexibility of catfish in coping with different environmental conditions, particularly in response to differences in prey abundnace and habitat characteristics between oligotrophic lakes and eutrophic reservoirs. Optimizing catfish activity strategies involves a dynamic interplay between evolutionary adaptations and behavioural flexibility that likely enhances the species’ ability to thrive in different ecological contexts. By understanding how catfish adapt to different environments, conservationists can better adapt management strategies to maintain and enhance or reduce catfish populations in different aquatic habitats. In addition, further analysis of the long-term consequences of these activity patterns, particularly in terms of growth rates and overall fitness, promises to provide deeper insights into the ecological dynamics of catfish populations in different waterbodies.

## Acknowledgements

Authors would like to acknowledge all FishEcU members for their help during fieldwork and data processing.

## Ethics statement

This study was approved by the Animal Welfare Committee of the Biology Centre CAS (45/2014) according to § 16a of the Act No. 246/1992 Coll., on the protection of animals against cruelty, and Experimental Animal Welfare Commission under the Ministry of Agriculture of the Czech Republic and with permission (Ref. no. 310/7387) of the managers of the study sites, Povodí Vltavy s.p. and DIAMO s.p..

## Data archiving statement

Data available from the github repository https://github.com/rubenrabb01/Hungry-Catfish-Effects-of-Prey-Availability-on-Movement-Dynamics-of-a-Top-Predator.git

## Author’s contributions

Milan Říha and Rubén Rabaneda-Bueno conceived the ideas and designed methodology; Milan Říha, Lukáš Vejřík, Marek Šmejkal, Martin Čech, Vladislav Draštík, Petr Blabolil, Michaela Holubová, Tomáš Jůza, Karl Ø. Gjelland, Zuzana Sajdlová, Luboš Kočvara, Michal Tušer and Jiří Peterka collected the data; Rubén Rabaneda-Bueno, Milan Říha, Lukáš Vejřík and Vladislav Draštík analysed the data; Milan Říha and Rubén Rabaneda-Bueno led the writing of the manuscript. All authors contributed critically to the drafts and gave final approval for publication.

