## Supplementary materials for "Hungry catfish – effect of prey availability on movement dynamics of a top predator"

### Appendix S1

#### Extended Methods

##### *Estimating the consistency of horizontal space utilisation (HSU)*

We analysed the consistency of horizontal space use across the study period by calculating the repeatability index (R), which represents the proportion of interindividual differences relative to the total variability of the behavioural response, which in mixed-effects models is the sum of variance components resulting from within- and between-individual differences. Both the agreement value and the adjusted R-value were calculated for each species and each diel period using separate GAM models. The agreement R-value was estimated using null models fitted without fixed effects and including only the random intercept (fishID) and the autocorrelation structure. Two adjusted R values were calculated, one for models with fixed effects of the covariates and the random intercept, and another value using intercept-slope models in which the slope of time was added to the individual random effects. In both cases, R is assumed to be constant for all values of the confounding variables (fixed and random). We calculated the R estimates for all models and inserted them into a table according to the following formula:

$$R = \frac{s(\tau_{00})}{s(\tau_{00}) + \emptyset}$$

where s(τ00) is the smooth random intercept and ϕ is the scale parameter, both estimated from the model (for more details see Jaric et. al, 2022).

*The parameterization of the GAMLSS model for habitat use*

GAMLSS are semi-parametric regression models that allow mixed distributions in which the parameters of the distribution can be modeled independently of each other with non-parametric functions (e.g. non-linear). Similar to the GAM models, the GAMLSS contain a scale parameter associated with the variance component, but they contain additional functions, each of which specifies the relationship between the predictors and one of the parameters, and additionally allow for mixed distributions other than the exponential family. GAMLSS models were fitted assuming a one-inflated beta distribution of the response variable, TOW, with three parameters, two location and scale parameters (μ , σ) and one shape parameter (v), modeled with the default link functions of the BEINF1 family, two logit links for the continuous values (0 < TOW < 1) of the beta-distributed variable BE(μ, σ) and a logit link for v to model the probabilities at 1 (p1). The fitted general model can thus be expressed as follows:

TOW ~ BEINF1(μ, σ, v)

$$g_1(\mu) = \eta_1 = X_1\beta_1 + \sum_{j=1}^{J_1} h_{j1}(x_{j1}) \quad (1.1)$$

$$g_2(\sigma) = \eta_2 = X_2\beta_2 + \sum_{j=1}^{J_2} h_{j2}(x_{j2}) \quad (1.2)$$

$$g_3(\nu) = \eta_3 = X_3\beta_3 + \sum_{j=1}^{J_3} h_{j3}(x_{j3}) \quad (1.3)$$

where μ, σ, v are the distribution parameter vectors of length $\eta_k$, with the link function $g_k$, with $\eta_1 = \text{logit}\left[\frac{\mu}{(1-\mu)}\right]$, $\eta_2 = \text{logit}\left[\frac{\sigma}{(1-\sigma)}\right]$ and $\eta_3 = \log(\nu)$; $\beta_k$ are parameter vectors of the linear (z-score) coefficients, which represent the proportion of partial use of open water ($TOW_p$), the proportion of variance in partial use of open water and the odds ratio of total use of open water ($TOW_t$), respectively, given in the response scale in the "gamlss" package; $X_k$ are design matrices for fixed parameter estimation containing the linear predictors; $h_{jk}$ are additive smoothing functions for fixed predictors that depend on random effects, such as $h_{jk}(x_{jk}) = h_{jk} = \Upsilon_{jk}$, where $\Upsilon_{jk}$ is the random additive term that includes the random (slope-intercept) effects and an ARMA autocorrelation structure of order (p=0, q=1).

.

The mixed continuous-discrete density function can be expressed as follows:

$$f_Y(\text{TOW}|\mu,\sigma,\nu) = \begin{cases} (1-p_0-p_1)\, f_W(\text{TOW}) & if\ 0 < \text{TOW} < 1\ (\text{TOW}_p) \\ p_1 & if\ \text{TOW} = 1\ \ (\text{TOW}_t) \end{cases}$$

(2)

for 0 ≤ y ≤ 1, where both $f_W(TOW) \sim BE(\mu,\sigma)$ with 0 < μ < 1, 0 < σ < 1, and the inflated beta with $\nu > 0$, and $p_1 = \frac{\nu}{1+\nu}$. The probability density function $f_W(TOW)$ is given by:

$$f_W(\text{TOW}) = \frac{1}{B(\alpha,\beta)} TOW^{\alpha-1}(1-TOW)^{\beta-1} \quad (3)$$

for 0 < TOW < 1, hence in (2),

$$\begin{pmatrix} \alpha \\ \beta \\ p_1 \end{pmatrix} = \begin{pmatrix} \mu(1-\sigma^2)/\sigma^2 \\ (1-\mu)(1-\sigma^2)/\sigma^2 \\ \nu/(1+\nu) \end{pmatrix} \tag{4}$$

with $\mathrm{E}(TOW) = \mu$ and $\mathrm{Var}(TOW) = \sigma^2\mu(1-\mu)$, and since ν is fitted with a log link function, it is defined as the odds ratio i.e., $\nu = \frac{p_1}{1-p_1}$ and $\nu = e^{\beta_k}$ in (1.3). Hence in (4),

$$\begin{pmatrix} \mu \\ \sigma \\ \nu \end{pmatrix} = \begin{pmatrix} \alpha(\alpha+\beta)^{-1} \\ (\alpha+\beta+1)^{-\frac{1}{2}} \\ \frac{p_1}{(1-`p_1)} \end{pmatrix} \tag{5}$$

Based on the above formulations, the final models fitted for each lake can be expressed with the following notations:

$$\{\mathrm{TOW}|OLIGOT1\} = \begin{bmatrix} \mathrm{logit}(\mu) = \alpha_0 + cs(\mathrm{time}) \times \mathrm{diel\_period} + bl + re(\sim\mathrm{time}|\mathrm{fishID}, \mathrm{corARMA}) \\ \mathrm{logit}(\sigma) = \beta_0 + bl + re(\sim\mathrm{time}|\mathrm{fishID}, \mathrm{corARMA}) \\ \log(\nu) = \delta_0 + \mathrm{diel\_period} + re(\sim\mathrm{time}|\mathrm{fishID}, \mathrm{corARMA}) \end{bmatrix}$$

$$\{\mathrm{TOW}|OLIGOT2\} = \begin{bmatrix} \mathrm{logit}(\mu) = \alpha_0 + cs(\mathrm{time}) + \mathrm{diel\_period} + pb(\mathrm{bl}) + re(\sim\mathrm{time}|\mathrm{fishID}, \mathrm{corARMA}) \\ \mathrm{logit}(\sigma) = \beta_0 + cs(\mathrm{time}) + \mathrm{diel\_period} + bl + re(\sim\mathrm{time}|\mathrm{fishID}, \mathrm{corARMA}) \\ \log(\nu) = \delta_0 + \mathrm{diel\_period} + re(\sim\mathrm{time}|\mathrm{fishID}, \mathrm{corARMA}) \end{bmatrix}$$

$$\{\mathrm{TOW}|EUT\} = \begin{bmatrix} \mathrm{logit}(\mu) = \alpha_0 + cs(\mathrm{time}) + \mathrm{diel\_period} + bl + re(\sim\mathrm{time}|\mathrm{fishID}, \mathrm{corARMA}) \\ \mathrm{logit}(\sigma) = \beta_0 + cs(\mathrm{time}) + \mathrm{diel\_period} + bl + re(\sim\mathrm{time}|\mathrm{fishID}, \mathrm{corARMA}) \\ \log(\nu) = \delta_0 + cs(\mathrm{time}) + \mathrm{diel\_period} + re(\sim\mathrm{time}|\mathrm{fishID}, \mathrm{corARMA}) \end{bmatrix}$$

### Figures

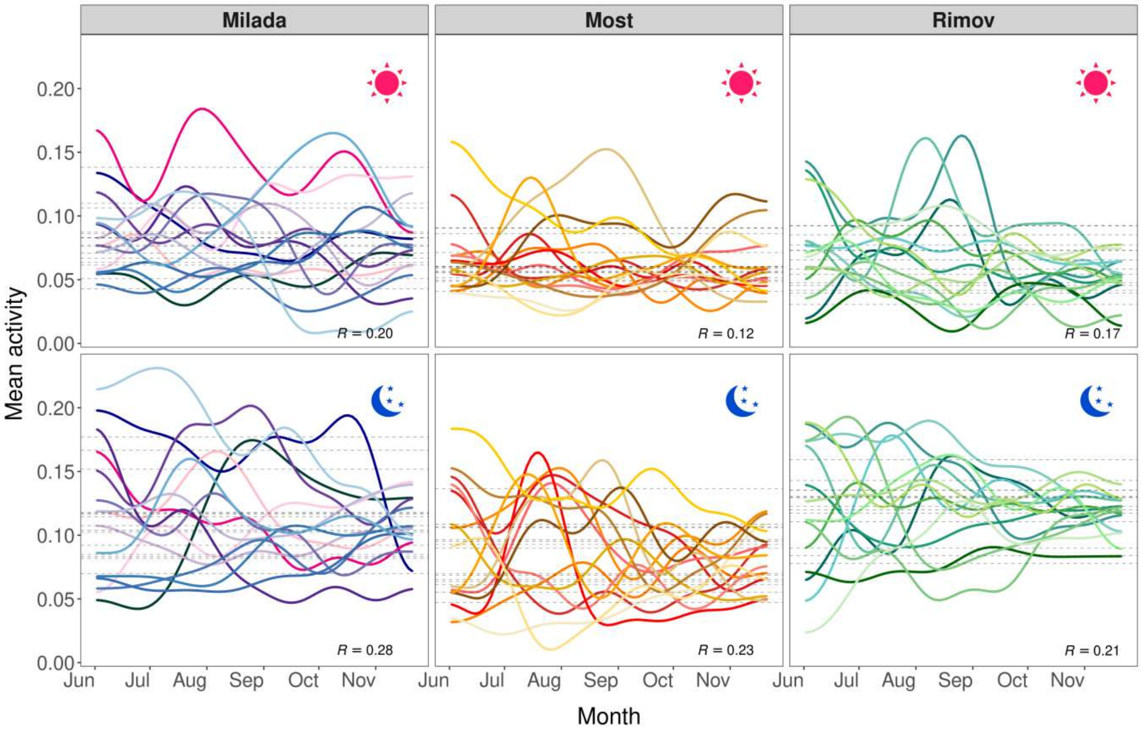


**Figure S1.** Individual trajectories of mean activity during the day (top) and night (bottom) estimated by GAM models. Presented are adjusted R-values from the final fitted model along with 95% confidence intervals derived using progressively complex GAMs. The modeling process initiates with random intercepts and incorporates fixed effects alongside random slopes.

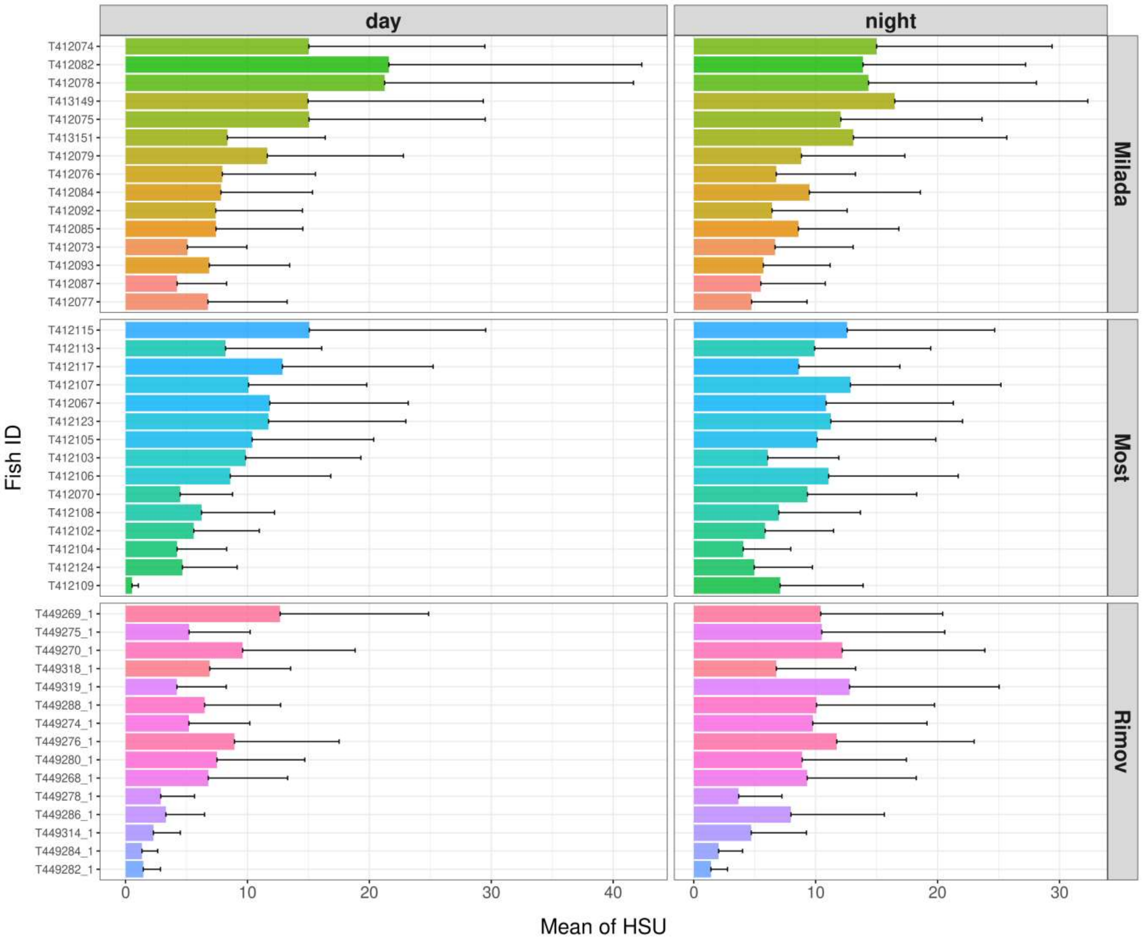


**Figure S2.** Boxplot depicting the mean habitat space use (HSU) along with error bars (standard deviation), estimated from GAM Models across three waterbodies.

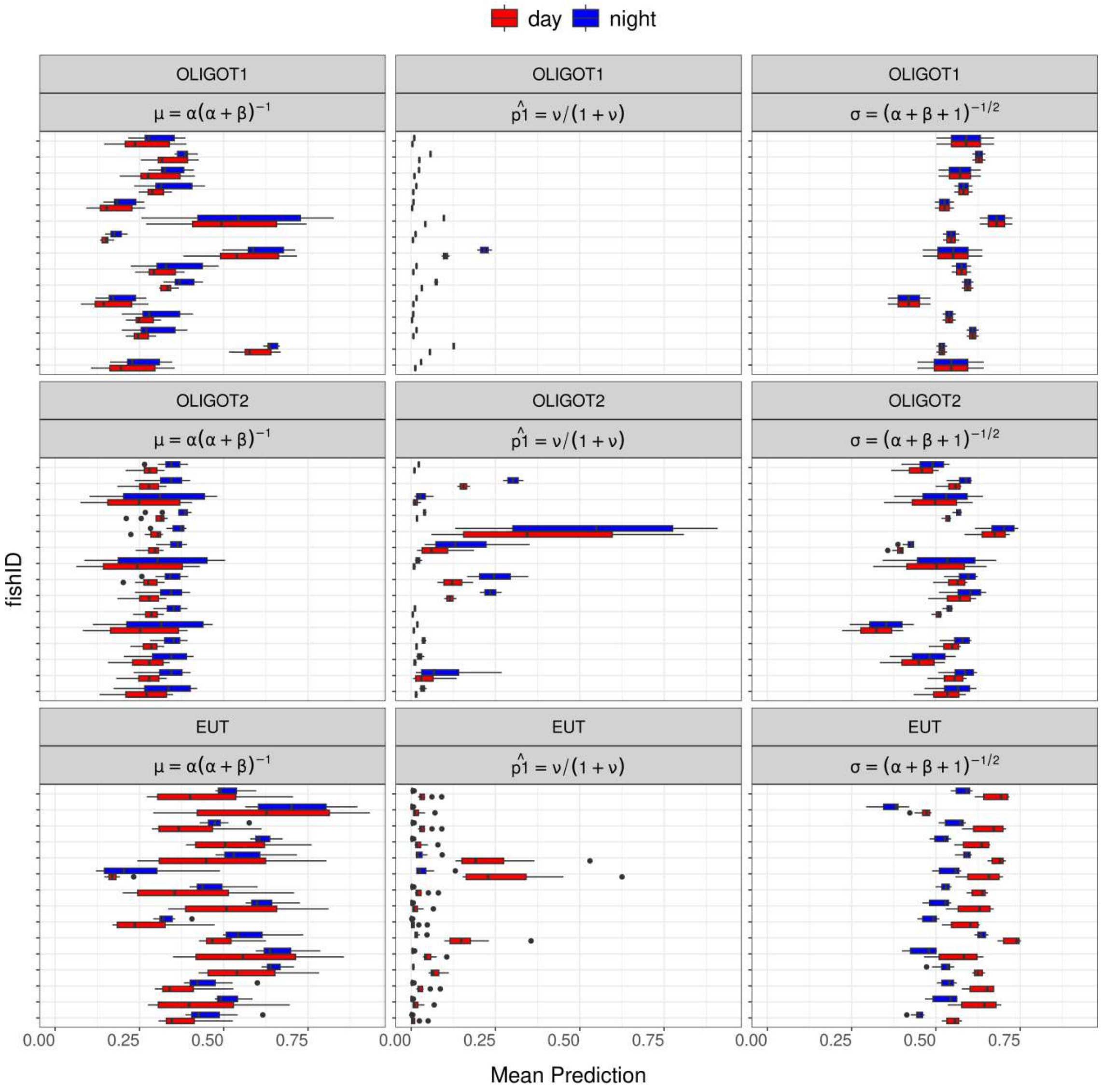


**Figure S3.** Predicted mean values of Time spent in Open Water (TOW) across waterbodies and diel periods using GAMLSS parameterization (see Extended Methods section in Appendix S1 for details). Each tick line on the y-axis represents an individual fish (fishID), with variations observed between waterbodies.

### Tables

**Table S1.** GAMLSS Models analyzing habitat use as a function of Time spent in Open Water (TOW). All models were fitted using a one-inflated beta distribution (BEINF1 in the R package “gamlss”) and are listed in order of best to worst fit based on their global fitted deviance (GDEV) and Schwarz Bayesian Criterion (SBC). 'df' indicates the degrees of freedom used for the three distribution parameters μ, σ, and v (in parenthesis), as well as the total number of degrees of freedom for the models. 't' denotes the continuous variable 'time' included as a fixed effect, while in the additive random term, 'time' was also included as a random slope alongside a random intercept (fishID); 'DP' represents the diel period factor variable; 'BL', body length; and 'pseudo-$R^2$', generalized pseudo R-squared (Nagelkerke 1991).

| Model | Description | df (μ) | df (σ) | df (v) | df | GDEV | SBC |
|---|---|---|---|---|---|---|---|
| | TOW\|OLIGOT1 ~ *BEINF1*(μ, σ, v) | | | | | | |
| **$Nu_1$** | **μ**($t^c$×DP, BL), σ(BL), **v**(DP) | -48.52 | 26.03 | 3.36 | -19.13 | -4191.4 | -4350.57 |
| **$Nu_2$** | **μ**($t^c$×DP, $BL^p$), σ(BL), **v**(DP) | -48.51 | 26.03 | 3.36 | -19.12 | -4191.39 | -4350.45 |
| **$Nu_3$** | **μ**($t^c$×DP, $BL^p$), σ($BL^p$), **v**(DP) | -48.5 | 26.04 | 3.36 | -19.11 | -4191.4 | -4350.37 |
| **$Nu_4$** | **μ**($t^p$×DP, $BL^p$), σ($BL^p$), **v**(DP) | -21.63 | 26.03 | 3.36 | 7.75 | -4306.49 | -4242.03 |
| **$Nu_5$** | **μ**($t^p$×DP, $BL^p$), σ($t^p$×DP, $BL^p$), **v**($t^p$×DP) | -21.2 | 31.88 | 22.99 | 33.66 | -4357.35 | -4077.31 |
| | TOW\|OLIGOT2 ~*BEINF1*(μ, σ, v) | | | | | | |
| **$Nu_1$** | **μ**($t^c$×DP, BS), σ($t^c$×DP, BS), **v**($t^c$, DP ) | 18.856 | 33.512 | 18.096 | 70.464 | -1040.216 | -483.118 |
| **$Nu_2$** | **μ**($t^c$, DP, $BL^p$), σ($t^c$, DP, $BL^p$), **v**(DP) | 18.856 | 33.515 | 18.096 | 70.467 | -1040.215 | -483.089 |
| **$Nu_3$** | **μ**($t^c$×DP, $BL^p$), σ($t^c$, DP, $BL^p$), **v**($t^c$, DP) | 19.437 | 33.512 | 26.333 | 79.282 | -1099.254 | -472.44 |

| | | | | | | | |
|---|---|---|---|---|---|---|---|
| **Nu$_4$** | **μ**($t^p$×DP), σ($t^p$, DP), **v**($t^p$, DP) | 23.798 | 28.878 | 22.802 | 75.478 | -1043.251 | -446.506 |
| **Nu$_5$** | **μ**($t^p$×DP, $BL^p$), σ($t^p$, DP), **v**($t^p$, DP) | 22.954 | 30.36 | 22.802 | 76.116 | -1044.513 | -442.727 |
| | TOW\|EUT ~*BEINF1*(μ, σ, v) | | | | | | |
| **Nu$_1$** | **μ**($t^c$×DP, BS), σ($t^c$×DP, BS), **v**($t^c$, DP ) | 19.790 | 30.577 | 18.647 | 69.015 | -1576.828 | -1014.127 |
| **Nu$_2$** | **μ**($t^c$×DP, $BL^p$), σ($t^c$×DP, $BL^p$), **v**($t^c$, DP) | 19.752 | 30.598 | 18.647 | 68.996 | -1576.605 | -1014.053 |
| **Nu$_3$** | **μ**($t^p$×DP, $BL^p$), σ($t^p$×DP, $BL^p$), **v**($t^p$, DP) | 35.807 | 35.878 | 24.046 | 95.731 | -1719.762 | -939.234 |
| **Nu$_4$** | **μ**($t^p$×DP, $BL^p$), σ($t^p$×DP, $BL^p$), **v**($t^p$×DP) | 35.807 | 35.878 | 25.118 | 96.803 | -1720.753 | -931.484 |
| **Nu$_5$** | **μ**($t^p$×DP, $BL^p$), σ($BL^p$), **v**(DP, $BL^p$) | 38.382 | 24.789 | 24.046 | 87.217 | -1546.432 | -835.324 |

$^p$ The predictor was fitted with a penalized P-spline smoothing function. Otherwise, the linear term is shown.
$^c$ The predictor was fitted with a cubic spline smoothing function. Otherwise, the linear term is shown.
